# Social Decision-Making in Response to Competing Stimuli

**DOI:** 10.64898/2026.10.05.756797

**Authors:** Emily K. Lessig, Hans A. Hofmann

**Affiliations:** Department of Integrative Biology, The University of Texas at Austin, Austin, TX, USA; Interdisciplinary Neuroscience Program, The University of Texas at Austin, Austin, TX, USA

**Author notes:** Correspondence to: Dr. Emily Lessig, Dr. Hans A. Hofmann, Department of Integrative Biology, The University of Texas at Austin, 2415 Speedway - C0930, Austin, TX 78745, USA. Rua da Quinta Grande, 6, 2780-156 Oeiras, Portugal: Dr. Emily Lessig, Gulbenkian Institute for Molecular Medicine, Integrative Behavioural Biology.

**Keywords:** social decision-making, Astatotilapia burtoni, dopamine, testosterone, social cognition, neural activity

## Abstract

Animals routinely encounter diverse social stimuli that can be simultaneous and vary in salience, valence and relevance in regard to ongoing behavioral decisions. Understanding how animals weigh such competing information is crucial for revealing the cognitive and neural mechanisms underlying context-dependent social behavior. Here, we investigated how male *Astatotilapia burtoni* navigate situations in which they must choose between rival males, potential mates, or both socially relevant cues. We first quantified focal male engagement toward stimulus types and found that males prioritized stimuli based on identity, context, and inferred threat level. We then examined the underlying neural correlates of these decisions in the preoptic area (POA) and area Dm (the putative teleost homolog of the basolateral amygdala (area Dm), two key nodes of the social decision-making network (SDMN). Using phosphorylated ribosomal protein S6 (pS6) as a marker of neural activity and tyrosine hydroxylase (TH) to identify dopaminergic cell populations, we found that overall activity of these brain regions as well as the activity of dopaminergic cell populations did not differ between the different social stimulus treatments. Our results reveal how animals navigate complex environments involving multiple social cues and highlight the behavioral and neural mechanisms supporting stimulus prioritization.

## INTRODUCTION

Decision-making is a fundamental attribute of animal behavior where individuals must decide what to respond to and what to ignore (Barnard 2005; Dill 1987). The salience of a stimulus, or the degree to which a stimulus attracts attention, can influence decision-making as animals may be more or less attuned to particular stimuli based on their sensory system, motivation, state, or experience. As animals often live in dynamic social environments where they are exposed to a multitude of stimuli simultaneously, context-dependent behavioral flexibility is necessary to raise Darwinian fitness (O’Connell and Hofmann 2011; Taborsky and Oliveira 2012). To make context-appropriate decisions (see “social competence” by Taborsky & Oliveira, 2012) animals have evolved cognitive mechanisms that can efficiently process and integrate different types of information (O’Connell and Hofmann 2011; Shettleworth 2010).

Behavioral responses to stimuli can therefore vary over time depending on a variety of internal as well as external factors such as available social information, environmental conditions, and other individuals (Jackson and Elwood 1989; Leger et al. 1979; Lima and Dill 1990). For example, the presence of an audience can influence foraging behavior (Leaver et al. 2007), male-male aggression (Cruz and Oliveira 2015; R. J. Matos and McGregor 2002; Roleira et al. 2017), as well as mating behavior (Plath et al. 2008). Desjardins et al., (2012) demonstrated that subordinate cichlid fish can display courtship and aggressive behavior when other subordinate males are watching but cease these behaviors when a dominant male is present. Similarly, bystanders can eavesdrop on agonistic displays and modify their behavior in response to the observed interactions (Abril-de-Abreu et al. 2015). Interestingly, a number of species have been found to use transitive inference or observe relationships to make inferences about unknown relationships (Davis 1992; Gillan 1981; Grosenick et al. 2007; von Fersen et al. 1991) which demonstrates social animals are keenly aware of their environment and modulate their behavior based on acquired social information.

In these dynamic social environments, animals face different types of opportunities (e.g. social ascension, mate) and challenges (e.g. predator, intruder). Many studies have examined how animals respond to challenges or opportunities including how sensory information influences behavioral, physiological, and neuronal responses (Alward et al. 2020; Maruska et al. 2013; Maruska and Fernald 2010; O’Connell et al. 2013). Field et al., (2018) demonstrated that males of Burton’s Mouthbrooder cichlid fish, *Astatotilapia burtoni*, need sexually relevant visual signals to engage in courtship behavior but increase this behavior when exposed to both visual and chemosensory signals compared to either alone. Further, Sessa et al., (2013) found that sex steroid hormones modulate behavioral responses to a social challenge and opportunity in male convict cichlids, *Amatitlania nigrofasciata*, where aggression was modulated by androgen receptor in a challenge context and digging was modulated by androgen and estrogen receptors in an opportunity context. However, in nature, observations show that males frequently encounter intruder challenges and reproductive opportunities simultaneously (Fernald and Hirata 1977; Desjardins et al. 2012), such that examining responses to a single stimulus may not capture the complexity of an animal’s naturalistic social environment (Barlow 2000). Do animals in this situation attend to both stimuli equally, and is one considered more salient than the other? How is this cognitive conflict represented in the brain? And which neuromodulators regulate responses to these competing stimuli? Surprisingly, few studies have investigated how animals deal with highly salient and competing stimuli. However, some behavioral work has examined how animals respond when multiple social or ecological cues are available simultaneously, including studies on competing displays in birds (Patricelli et al. 2002), mate choice under simultaneous options in fish (Jeswiet and Godin 2011), and threat-opportunity tradeoffs in territorial species (Ord and Evans 2002; Candolin 2004). These examples illustrate that animals can shift attention dynamically when confronted with concurrent demands, but the underlying mechanisms remain poorly understood.

Acute endocrine responses are also important in both challenge and opportunity contexts, perhaps modulating subsequent behavior. The “challenge hypothesis” was originally proposed to explain seasonal variations in androgen levels, such as testosterone, in relation to competition and parental care in birds (Wingfield et al. 1990) but similar patterns have been demonstrated across vertebrates, demonstrating that males of diverse species exhibit elevated levels of testosterone in response to social challenges (Hirschenhauser and Oliveira 2006). However, it has long been known that access to females can also elevate androgen levels in males (Rose et al. 1972; O’Connell et al. 2013), especially in males who do not exhibit parental care. For example, the role of receptive females can have both an indirect effect on male androgen levels because competition among males is strongest when fertile females are available, but also a direct effect via male-female interactions (Amstislavskaya and Popova 2004; Goymann et al. 2007; Hirschenhauser and Oliveira 2006; Wingfield et al. 1990).

Androgens can also regulate behavior by binding to nuclear receptors that act as transcription factors in the brain. Many brain regions that are responsive to sex steroid hormones are part of the Social Decision-Making Network (SDMN), an evolutionarily ancient neural circuit involved in the integration of social stimuli and reward processing across vertebrates (O’Connell and Hofmann 2011; 2012). Previous work on the neural basis of social interactions suggests several candidate brains regions that are of particular importance in studies on social decision-making, especially in the context of aggression and reproduction. For example, the preoptic area (POA) mediates sexual and aggressive behavior across vertebrates (Hull and Dominguez 2006) and the basolateral amygdala is involved in valence coding and mediating aggressive behavior (Pignatelli and Beyeler 2019).

Along with steroid hormones, the SDMN involves other neuroendocrine and neuromodulatory pathways such as the monoaminergic system (including serotonin, dopamine, and noradrenaline) and is highly sensitive to dopaminergic mediation (O’Connell and Hofmann 2011; 2012). Dopamine (DA) is a neurotransmitter that modulates different animal behaviors such as evaluating risk, encoding the rewarding properties of a stimulus and anticipating the outcomes of social interactions (Berridge and Robinson 1998; Heimovics et al. 2009), which supports context-appropriate decision-making (Schultz 1998, 2002). DA also plays an important role in motivation, learning and memory (Dalley and Everitt 2009; Flagel et al. 2011; Goldman-Rakic 1997; Krashes et al. 2009; Wise 2004) functions supported by brain regions such as the hippocampus, amygdala, and preoptic area in mammals and contribute to the vertebrate mesolimbic reward system (Fried et al. 2001; Kempadoo et al. 2016; McNamara et al. 2014; Rosenkranz and Grace 1999). In teleost fish, dopaminergic circuits are similarly central to social behavior and experience-dependent plasticity. For example, changes in social status rapidly alter dopaminergic receptor expression and synthesis pathways in *Astatotilapia burtoni* (Loveland et al. 2014; Maruska et al. 2013), and social interactions modulate DA levels in other teleost such as zebrafish, *Danio rerio* (Teles et al. 2013; Polavieja and Orger 2018). Additionally, DA neurons in fish encode prediction and support associative learning (Dreosti et al. 2015; Antunes et al. 2022), highlighting a conserved role for dopamine in motivating and shaping social behavior across vertebrates.

DA also reinforces sexual behavior such as pair bonding in the monogamous prairie vole (*Microtus ochrogaster*) and mating behavior in rats (*Rattus norvegicus*) (Curtis et al. 2006; Dominguez and Hull 2005; Hull and Dominguez 2006). Additionally, DA mediates aggressive behavior such as in Arctic charr (*Salvelinus alpinus*) where subordinate males have lower dopaminergic activity and lower aggression (Winberg et al. 1991), and in the African cichlid fish (*Astatotilapia burtoni*) pharmacological blockage of DA receptor reduces male aggression depending on social context (Weitekamp et al. 2017). The dopaminergic reward system is therefore of particular interest in understanding how animals evaluate the salience of social stimuli, especially intruder challenges and reproductive opportunities.

We used Burton’s Mouthbrooder cichlid, *Astatotilapia burtoni*, to investigate how males respond to competing stimuli. *A. burtoni* males have two distinct and reversible phenotypes: dominant (10-30% of population) and subordinate (70-90% of population). Dominant males are brightly colored, aggressively defend spawning territories, and reproduce with females. Subordinate males are drably colored, do not hold territories, and are reproductively suppressed (Fernald and Hirata 1977; Huffman et al. 2015; Maruska and Fernald 2010). *A. burtoni* is an ideal system to study social cognition as males display context-dependent behavior that can be robustly quantified and manipulated in naturalistic laboratory settings (Hofmann 2003; Karen P. Maruska and Fernald 2018). For example, *A. burtoni* males modify their behavior both by observing other individuals (e.g. transitive inference: Grosenick et al., 2007) as well as being observed by other individuals (e.g. audience effect: Desjardins et al., 2012). Additionally, males mediate their aggression based on familiarity with other individuals, reducing aggression to familiar individuals. Once familiar, *A. burtoni* males can also cooperate in territorial defense and jointly aggress towards intruder males (Weitekamp et al. 2017).

Importantly, the SDMN and dopaminergic system have been described in detail in this species, suggesting candidate brain regions that are of importance in social behavior and decision-making (O’Connell et al. 2013; O’Connell and Hofmann 2011). In particular, the mammalian preoptic area (POA) is involved in sexual and aggressive behavior (Hull and Dominguez 2006) and has a putative homolog in teleost fish (POA: (O’Connell and Hofmann 2011); the mammalian striatum is in involved information and reward processing (Báez-Mendoza and Schultz 2013; Balleine et al. 2007) and has a putative homolog in teleost fish (area Vc: O’Connell & Hofmann, 2011); the ventral pallial component of the mammalian amygdala (BLA) encodes emotional and avoidance behavior (Adolphs 2010; Pignatelli and Beyeler 2019) and has a putative homolog in teleost fish (Dm: Mueller et al., 2011; Portavella et al., 2004; Rodríguez et al., 2002).

In the present study, we used naturalistic *A. burtoni* communities to test how males respond to both an intruder challenge and reproductive opportunity. Specifically, we exposed focal males to either intruder males, reproductive females, or both simultaneously and measured their behavioral responses. Because we were also interested in how hormonal and neural responses contribute to context-appropriate behavior, we quantified neural activity marker pS6 in brain regions known to be involved in sexual and aggressive behavior. We also examined the role of the dopaminergic system by labeling tyrosine hydroxylase (TH), the rate-limiting enzyme in catecholamine synthesis and a reliable neurochemical marker for dopaminergic cell populations (O’Connell et al. 2011). We hypothesized that males would display stimulus-specific behavioral responses-aggressive displays towards intruders and courtships displays towards reproductive females. We further hypothesized that males will also differ in their physiological and neural responses in key nodes of the SDMN in a stimulus-specific way.

## MATERIALS AND METHODS

### Animals

*Astatotilapia burtoni* descended from a wild-caught stock population were kept in 100 l aquaria under conditions mimicking their natural environment. Fish were fed daily with cichlid flakes (Omega One Cichlid Flake Fish Food, Arcata Pet Supplies, Arcata, CA) and a 12:12 h light: dark cycle was maintained. All work was approved by the Institutional Animal Care and Use Committee at the University of Texas at Austin.

### Behavior Trials

After males had been dominant for at least 1 week in a naturalistic community, they were placed into 100 l experimental tanks for a 7-day acclimation period where they were housed in compartments with 2 juveniles and 2 females each (Fig. 1). On night 7, stimulus fish were placed in the experimental tank to acclimate overnight, separated from the focal male by opaque acrylic dividers. On the morning of day 8, the opaque dividers were replaced with transparent ones and males were exposed to different social stimuli for 1 h (between 8:30-10:30h) in the intruder treatment (n=12), female treatment (n=12), or competing stimuli treatment (n=12) (Fig. 1). Stimulus fish were randomly assigned to either side compartment and all males were size-matched. Behavior was recorded for 1 h using a digital video surveillance system (Alibi). We scored the behavior of each focal male for 10min (10:20-10:30h) using the BORIS event recoding software (Friard and Gamba 2016) including aggressive displays (lateral display, border conflict, frontal display), courtship displays (leads and quivers), interaction time (s), as well latency for stimulus fish to appear (s) and focal male to approach them (s) (Supplementary Table 1). Interaction time was measured as the time both the focal male and stimulus fish spent within 1inch of the clear divider during the 10min observation window. The standard length (mm) and body mass (g) were measured, the gonadosomatic index (GSI) was calculated ((gonad weight (g) / body mass (g)) × 100), and body color was recorded for each focal male. Sizes ranged from 47mm – 60mm and 2.24g – 4.90g with GSI ranging from 0.04 – 1.02.

**Figure 1:**
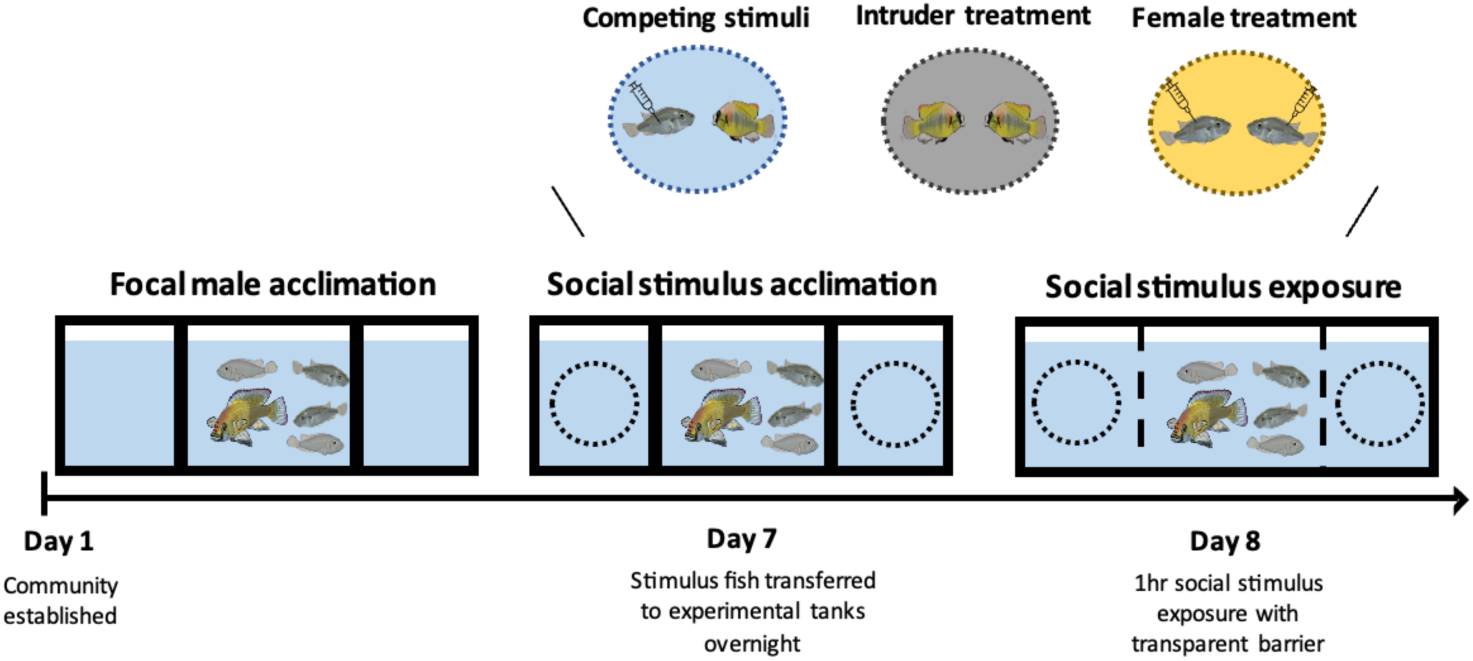
Experimental paradigm. Dominant *A. burtoni* males were given a 7-day acclimation period in experimental tanks with 2 juveniles and 2 females. Stimulus fish were added to the experimental tank on night 7-either 2 gravid females (female treatment), 2 intruder males (intruder treatment) or both a female and intruder (competing stimuli)-separated by opaque acrylic dividers. On day 8, focal males were given visual exposure to social stimuli through clear acrylic dividers for 1 h.

#### Intruder treatment

Intruder males were confirmed to be dominant for at least 1 week in a naturalistic community prior to being used for testing. The night before testing, a size-matched (between 9x% and 10x% of focal male SL) intruder male was placed on either side of the focal male’s compartment (Fig. 1). At the start of experimental period, focal males were given visual exposure to intruder males through clear acrylic dividers for 1 h.

#### Reproductive opportunity treatment

On the evening before testing, a gravid (egg bearing) female was placed on either side of the focal male’s compartment. Females were confirmed to be gravid upon visual inspection. At least 15 min before the start of the experimental period, females received 6ml injections of prostaglandin F_2_α (PGF_2_α; catalogue number 16010; Cayman Chemicals) via syringe intraperitoneally, diluted to 0.4% with 1X PBS. PGF_2_α has been shown to induce reproductive behavior (Juntti et al. 2016; Kidd et al. 2010b). At the start of the stimulus period, focal males were given visual exposure to receptive females through clear acrylic dividers for 1 h.

#### Competing stimuli treatment

In the competing stimuli paradigm, a gravid female was placed in one side compartment, and an intruder male was placed in the other side compartment next to the focal male on the evening before testing. Prior to the start of the stimulus period, females were injected with PGF_2_α as described above. Focal males were then given visual exposure to both the gravid female and intruder male through clear acrylic dividers for 1 h.

### Tissue collection

Following the 1 h stimulus exposure, focal males were removed from experimental tanks, measured and weighed, and then blood was collected through the dorsal aorta using heparinized 26-G butterfly infusion sets (SURFLO Winged Infusion Sets, Terumo). Focal males were then killed by rapid cervical transection and brains rapidly dissected. One hour is an appropriate time course for measuring neuronal activity by quantification of phosphorylated ribosomal protein S6 (pS6), a ribosome that becomes phosphorylated when neurons are actively translated (Knight et al. 2012; Ruvinsky and Meyuhas 2006). Brains were fixed overnight in 4% paraformaldehyde at 4°C, then washed in 1 X phosphate-buffered saline (PBS) and cryo-protected overnight in 30% sucrose at 4°C and finally embedded in O.C.T. compound (Tissue-Tek; Fisher Scientific Co., Pittsburgh, PA, USA). Brains and plasma were stored at -80°C until further processing.

### Hormone assay

Blood was spun down in a centrifuge at 3500rpm for 15min. Plasma was then separated and stored at -80°C. Plasma testosterone was quantified using a competitive assay (Cayman Chemicals, Ann Arbor, MI, USA) in accordance with the manufacturer’s instructions. While 11-ketotestosterone (11-KT) is the major androgen controlling male reproduction in many teleost fish (Borg 1994), circulating levels of 11-KT are much lower and more variable than those of testosterone in haplochromine cichlids including *A. burtoni* (Dijkstra et al. 2012; Kidd et al. 2010). In our study, the average intra-assay coefficient of variation (CV) was 6.28%.

### Immunohistochemistry (IHC)

Brains were sectioned on a cryostat at 30μm and thaw-mounted on Super-Frost Plus slides (Fisher Scientific) in three series that were stored at -80°C until further processing. We used immunohistochemistry (IHC) to visualize phosphorylated ribosomal protein S6 (pS6), a structural component of the ribosome that becomes phosphorylated when neurons are activated (Knight et al. 2012; Ruvinsky and Meyuhas 2006) and tyrosine hydroxylase (TH), the rate-limiting enzyme in catecholamine synthesis and a reliable neurochemical marker for dopaminergic cell populations in *A. burtoni* (O’Connell et al. 2011). An alternate series was used for pS6 and TH fluorescent double-labeling. Sections were removed from -80°C, dried on a slide warmer, and processed for IHC as has been previously described (Weitekamp and Hofmann 2017). Briefly, slides were washed 3 times in 1 x PBS and incubated in a blocking solution of 5% normal goat serum (NGS), 0.3% Triton X-100 and 1 x PBS for 2 hours at room temperature. Slides were then incubated in a mix of 2% NGS, 0.3% Triton X-100, 1:500 rabbit anti-pS6 primary antibody (Cell Signaling), 1:500 mouse anti-TH primary antibody (Vector Laboratories), and 1 x PBS overnight at room temperature. Slides were washed 3 times in 1 x PBS and incubated for 2h in a mix of 1:350 goat anti-rabbit Alexa Fluor 594 and 1:350 goat anti-mouse Alexa Fluor 488. Slides were then washed in 1 x PBS and cover-slipped with Vectashield hardset mounting media with DAPI (Vector Laboratories, Burlingame, CA, USA).

### Quantification of activated neurons

We quantified pS6 and TH staining in brain regions implicated in social behavior (putative homologs according to (O’Connell and Hofmann 2011): the medial part of the dorsal telencephalon (area Dm: the putative homolog of the basolateral amygdala) and the preoptic area (POA). For each brain region and individual, pS6 and TH-positive cells were counted across three sections using the Imaris software (v.10.2; Oxford Instruments). Briefly, a region of interest was defined using the ‘surfaces’ tool and positively labeled cells were counted (at 20x objective) using the ‘spots’ tool. Co-labeled cells were identified using ‘object-object statistics’ to quantify pS6 and TH labeled cells that were overlapping. For each brain region, pS6-labeled and pS6/TH colabeled cells were quantified across three sections. Cell counts were normalized to the quantified area and adjusted for section thickness (30*μ*m) to generate a standardized measure of cell abundance.

### Data Analysis

All analyses were conducted in R v. 4.4.2 with a significance threshold of *p* < 0.05. For behavior data, lateral displays, border conflicts and frontal displays were summed into aggressive displays and leads and quivers were summed into courtships displays. Interaction time was summed for the 10min observational period. To compare how focal males engaged with same sex stimuli (either intruder males or reproductive females), we conducted a paired samples t-test, where engagement for each focal male was separated into high vs. low. Welch’s t-test was conducted to compare whether focal males engaged more with intruder males or reproductive females in the control treatments, where only the high engagement scores were used for each focal male. We conducted an ANOVA to compare engagement and interaction time across all treatments, including competing sex stimuli. For *post-hoc* analyses, Tukey’s honestly significant difference (HSD) test was used. To test whether there were any differences in testosterone measures across treatments, we conducted an ANOVA with a Tukey HSD *post-hoc* analysis. To assess differences in neural activity across stimulus treatments, we conducted one-way ANOVAs for the POA and Dm comparing pS6-labeled cells and pS6/TH colabeled cells across treatments.

## RESULTS

### Dominant *A. burtoni* males preferentially engage with one stimulus animal when simultaneously exposed to two conspecifics of the same sex

We first examined whether males prefer to engage with one stimulus animal more than another, even if they are the same stimulus types (intruder male or reproductive female, respectively). We found that focal males engaged significantly more with one stimulus animal compared to the other, even when they are of the same sex (Fig. 2A & B; t = 4.475, df = 11, *p* = 0.0009; t = 5.824; df = 11; *p* = 0.0001). Further, in the absence of a competing sex stimulus, focal males engaged more with reproductive females compared to intruder males (Fig. 2C; t = 2.474, df = 15.61, *p* = 0.025). Interestingly, however, we did not find a difference in interaction time in the same sex treatments, where focal males did not spend a significantly different amount of time with either intruder male (*p* = 0.975) or reproductive female (*p* = 0.923) when presented simultaneously.

**Figure 2.**
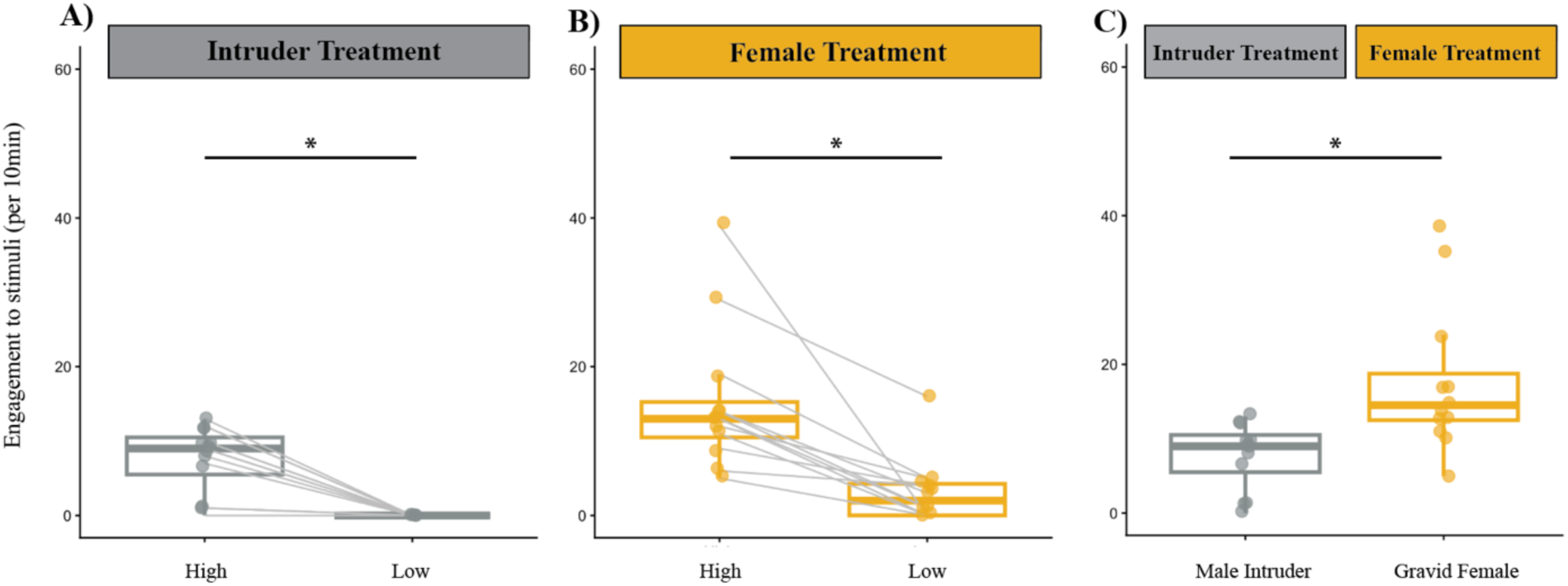
Focal male behavior in response to same sex stimuli. (A) In the intruder treatment, high vs low engagement is shown separately. (B) In the female treatment, engagement broken up by high vs low. (C) The female control and intruder control data combined where only ‘high’ levels of engagement are used for each focal male.

### Dominant *A. burtoni* males dynamically adjust their behavior when exposed to competing opposite sex stimulus

Next, we asked how males engage with both a challenge and reproductive opportunity when presented simultaneously. We found a significant difference in engagement based on social stimulus (ANOVA: F_3,44_=6.051, *p* = 0.001; Fig. 3A). When faced with competing opposite sex stimuli, focal males engaged more with male intruders compared to reproductive females (*p* = 0.004) or males in the intruder only treatment (*p* = 0.002). However, focal males did not engage more with reproductive females in the competing stimuli treatment compared to the female only treatment (*p* = 0.667). Additionally, males spent a different amount of time interacting with a challenge and opportunity when they were presented simultaneously (ANOVA: F_5,66_=3.441, *p* = 0.008; Fig. 3B). When faced with competing opposite sex stimuli, we found that focal males spent more time interacting with intruder males compared to reproductive females (*p* = 0.042). We also found that males in the competing stimuli treatment spent more time with intruder males compared to either gravid female in the female only treatment (*p* = 0.007 & *p* = 0.025; Fig. 3B).

**Figure 3.**
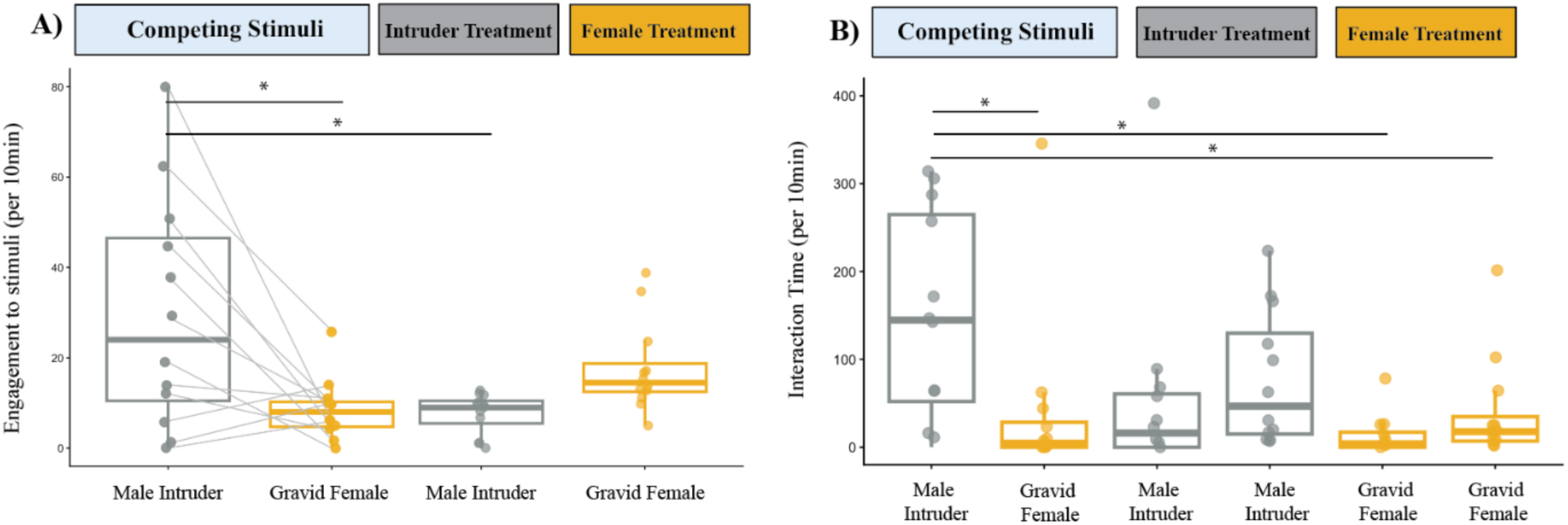
Focal male engagement across treatments. (A) Focal male behavior across all treatments. In the competing stimuli treatment, males engage more with the male intruder compared to the gravid female. Males also engage more intruders in the competing stimuli treatment compared to the intruder control. In intruder and female treatments, engagement is broken up by high vs. low. (B) Focal male interaction time across all treatments.

### Dominant males increase testosterone levels in response to reproductive opportunity

To understand the physiological responses of dominant males to both an intruder challenge and reproductive opportunity, we examined circulating testosterone levels of focal males following social stimulus exposure. As expected, circulating testosterone levels were elevated in all treatments compared to previously published baseline levels (Parikh et al. 2006; Winton et al. 2012). We found that circulating levels of testosterone increased even further when focal males were exposed to reproductive females in the female only treatment (ANOVA: F_2,33_ = 7.48, *p* = 0.0021; Fig. 4). However, testosterone levels in any of the treatments where an intruder male was present did not vary significantly from each other (*p* = 0.5041). We next tested whether testosterone levels were correlated with behavioral metrics. There were no significant correlations between testosterone levels and total engagement (R = -0.21, *p* = 0.079).

**Figure 4.**
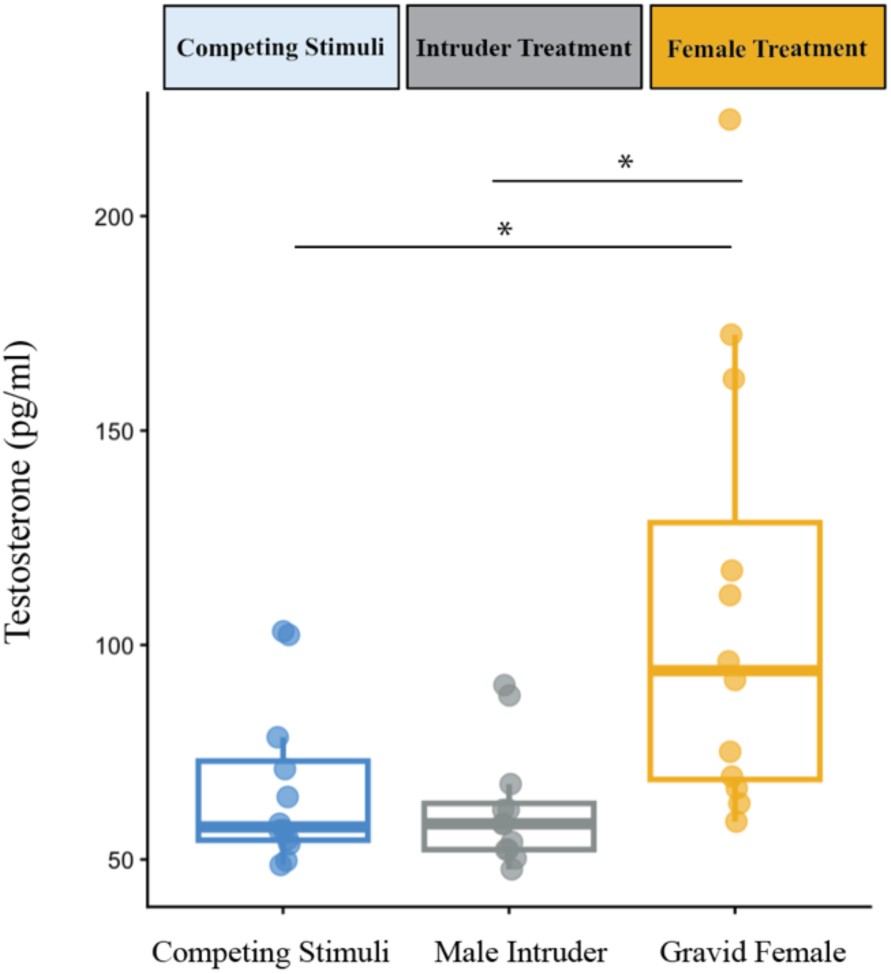
Focal male hormonal response to social stimuli. Males had higher levels of plasma testosterone in the female control compared to the other experimental treatments. Asterisks indicate significant differences (*p* < 0.05).

### Dominant males have similar levels of neural activity in response to different stimulus types

We next compared neural activity in the POA and area Dm in response to a male intruder, a gravid female, or both stimuli presented simultaneously. We found no significant differences among stimulus treatments in the number of pS6 and TH-colabeled cells in either the POA (ANOVA: F_2,18_=0.487, *p* = 0.622; Fig. 5A) or Dm (F_2,19_=0.767, *p* = 0.478; Fig. 5B). Similarly, the number of pS6-labeled cells did not differ among treatments in the POA (F_2,18_=0.363, *p* = 0.700; Fig. 5C) or Dm (F_2,20_=1.035, *p* = 0.373; Fig. 5D). Moreover, pS6 and TH-colabeling in the POA and area Dm was not significantly correlated with either total behavioral engagement with the social stimuli or circulating testosterone levels (Supplemental Figs. 2-3). Thus, neural activity in these regions, including activity within TH-positive cells, was similar whether males were exposed to competing male and female stimuli, a male intruder alone, or a gravid female alone.

**Figure 5.**
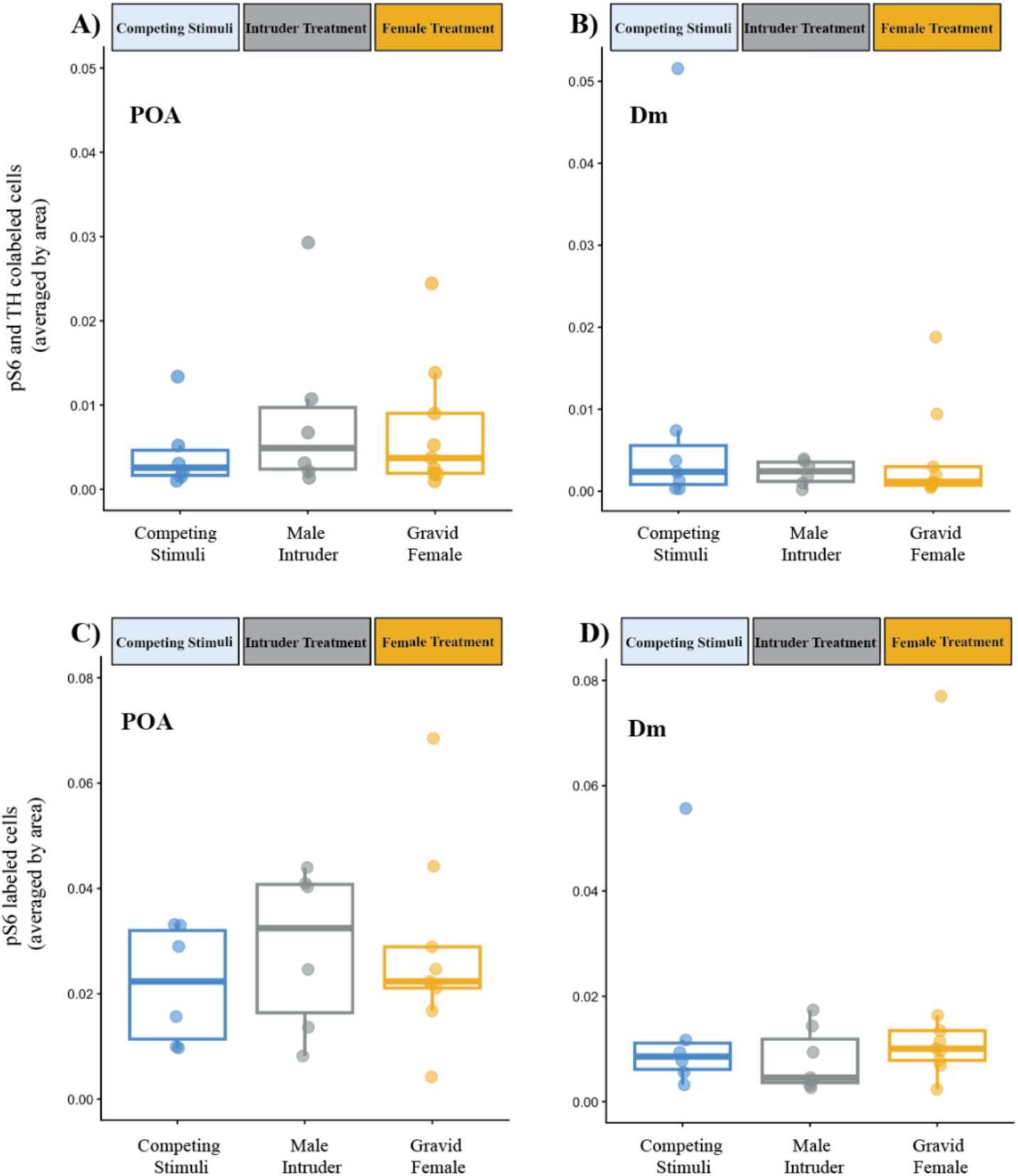
Neural activity in response to different stimulus types. Dominant males show similar levels of pS6 and TH colabeled cells in the POA (A) and Dm (B) as well as pS6 labeled cells (C&D).

## DISCUSSION

Animals living in dynamic social environments frequently encounter multiple stimuli at once, forcing them to prioritize, ignore, or integrate competing sources of information. In the present study, we examined how dominant *A. burtoni* males respond both behaviorally and physiologically to competing social stimuli: a male intruder, a gravid female, or both presented simultaneously. We found that males differentiate stimulus types through distinct behavioral patterns—displaying higher levels of aggression toward male intruders and increased courtship toward gravid females. Continued work will help us understand how animals are integrating multiple social cues in key brain regions in the Social Decision-Making Network (SDMN) to coordinate context-dependent behavior.

### Behavioral responses indicate stimulus prioritization but also behavioral flexibility

As predicted, males displayed aggressive behavior toward intruder males and courtship behavior toward gravid females, consistent with previous work showing stimulus-dependent behavior in *A. burtoni* (Maruska and Fernald 2010). When females and intruders were presented simultaneously, males still expressed both behaviors, demonstrating that they do not exclusively commit to one behavioral response but maintain behavioral flexibility, a key component of social competence (Taborsky and Oliveira 2012). However, males reduced behavior directed towards females and increased intruder-directed behaviors during the simultaneous treatments. This could indicate that males evaluate the relative salience of each stimulus and redistribute effort accordingly, a pattern consistent with theoretical predictions of salience-based decision-making. The elevated aggression toward intruders in the presence of a gravid female may also reflect an audience effect, in which males intensify competitive signaling when potential mates are observing, a phenomenon documented in other fish (Matos and McGregor 2002; Cruz and Oliveira 2015; Desjardins et al. 2012). Similar audience effects have been described across vertebrates where males amplify courtship or competitive displays when observed by potential mates or rivals (Peake et al. 2005; Earley 2010). For example, in field crickets, males increase signaling effort and aggression when females are present, even if the female is not the direct target of the interaction, indicating that males adjust investment based on the broader social audience rather than just the immediate opponent (Fitzsimmons and Bertram 2013; Montroy et al. 2016). These patterns reinforce the idea that male aggressive signaling is shaped not only by intruder threat but by the broader social context in which threats occur. Additional work further examining how the strength of each stimulus influences decision-making could help determine how males prioritize or weigh stimulus-based information leading to behavioral outputs.

### Neurophysiological responses reflect broad functional roles of POA and Dm

We quantified neural activity using ps6 and TH co-labeling in two nodes of the Social Decision-Making Network: the preoptic area (POA) and the dorsomedial telencephalon (Dm). Both regions are well-established mediators of reproductive behavior, aggression, and social valence across vertebrates (Hull and Dominguez 2007; Pignatelli and Beyeler 2019; O’Connell and Hofmann 2011). Given these broad functional roles, it is maybe not surprising that neural activity did not clearly differ among treatments as males often exhibited both aggressive and reproductive behavior. This functional overlap also extends to endocrine responses: testosterone modulates both aggressive and reproductive behavior and its secretion can be rapid and transient (Gleason et al. 2009; Marler et al. 2005) therefore highlighting that underlying physiological systems may exhibit overlapping activity, reflecting shared roles in supporting different dimensions of social behavior. Additional work examining the neural activity in these brain regions of interest will help to clarify these current results.

## CONCLUSIONS

This study examines how dominant *A. burtoni* males navigate competing social demands, that requires the integration of different forms of competing information to respond in real-time. Males responded behaviorally in a stimulus-specific manner, escalating aggression towards intruders and directing courtship behavior toward gravid females, yet flexibly redistributing effort when both stimuli were present. Neurophysiological responses in the POA and Dm did not differ across social contexts, while circulating testosterone was elevated in response to female stimuli suggesting that behavioral prioritization of competing social stimuli was not accompanied by corresponding treatment-specific differences in activity within these brain regions. These results highlight the importance of understanding decision-making in response to naturalistic and complex environments rather than single-stimulus context, as such complexity reveals how social competence emerges from integration of behavioral, hormonal, and neural processes. Future work expanding upon these results and incorporating additional SDMN nodes will further our understanding of how vertebrate brains resolve the tradeoffs imposed by competing social stimuli.

## Acknowledgements

We thank members of the Hofmann lab for discussion and Mike Ryan and Felicity Muth for commenting on earlier versions of the manuscript. We thank Jennie DeVore for technical assistance and Camille Brown for animal care. This work was supported by a Stengl-Wyer Fellowship, American Association of University Women Fellowship, Animal Behavior Society Student Research Grant (E.K.L.) and by NSF grant IOS-2444903 and the Reeder Endowed Fellowship in Evolutionary Biology (H.A.H.). Computational analyses were performed using the Biomedical Research Computing Facility at UT Austin, Center for Biomedical Research Support (RRID#: SCR_021979).

