## Supplementary materials for "Social Decision-Making in Response to Competing Stimuli"

**Supplementary Material**

**Supplemental Table 1. Behavioral displays quantified for focal males**

| **Type** | **Display** | **Description** |
| --- | --- | --- |
| **Aggressive** | Frontal display | One male aggressively displays towards another |
|  | Lateral display | Aggressive display where male presents his fins laterally with occasional quivering |
|  | Border conflict | Confrontation between two males to define territorial boundaries |
| **Reproductive** | Approach/Lead | Males swim towards females |
|  | Quiver | Male approaches female and presents fins laterally, occasionally quivering |
| **Social** | Interaction time | When both focal male and stimulus fish are interacting within ~1inch of clear divider |

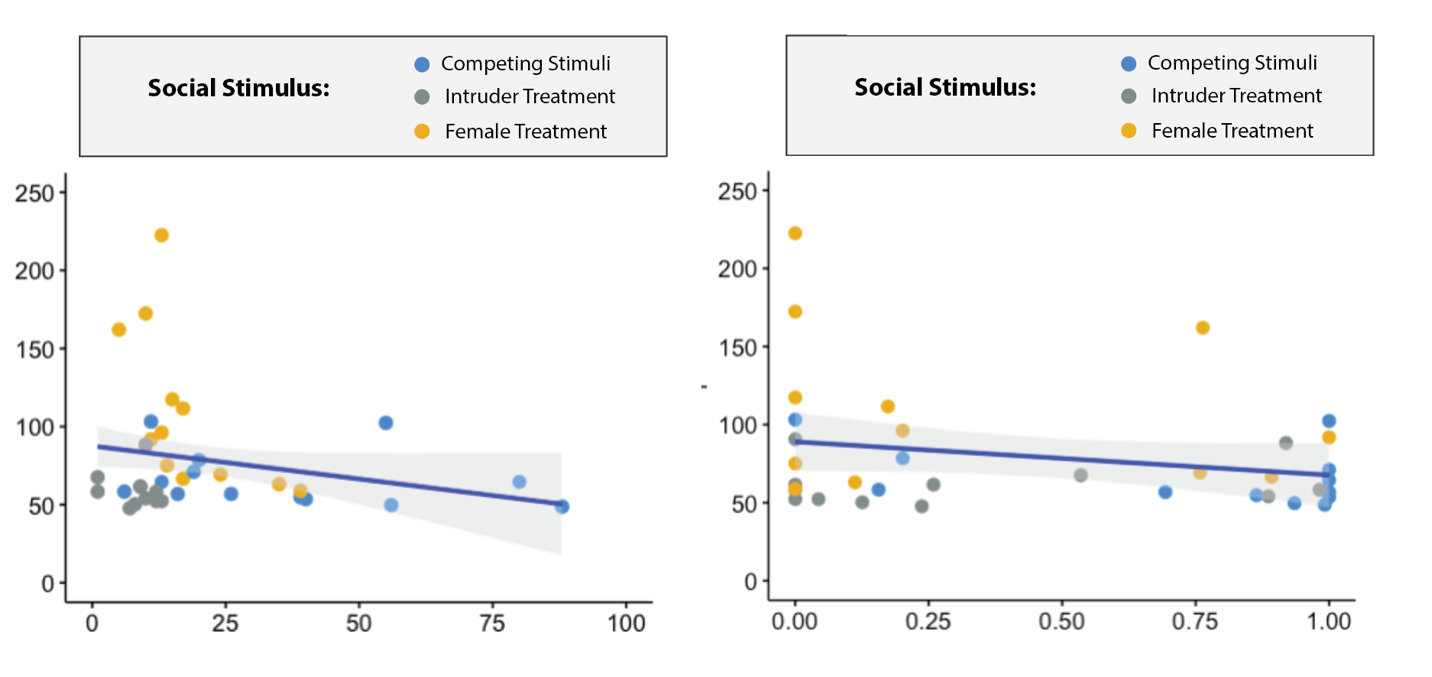

Preference Score

Engagement (total) to stimuli (per 10min)

Testosterone (pg/ml)

**Supplemental Figure 1. Relationship between circulating testosterone and behavioral engagement with social stimuli.** Circulating testosterone levels plotted against (A) total engagement with social stimuli during observational period and (B) preference score or proportion of total engagement directed toward the ‘preferred’ stimulus. Points represent individual focal males and are colored by social stimulus treatment. Lines represent linear regression fits with shaded 95% confidence intervals.

**B)**

**A)**

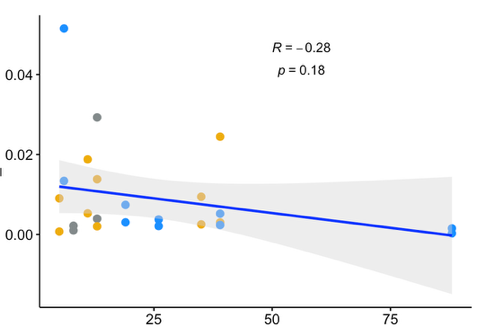

pS6 and TH colabeled cells (averaged by area)

**DM & POA**

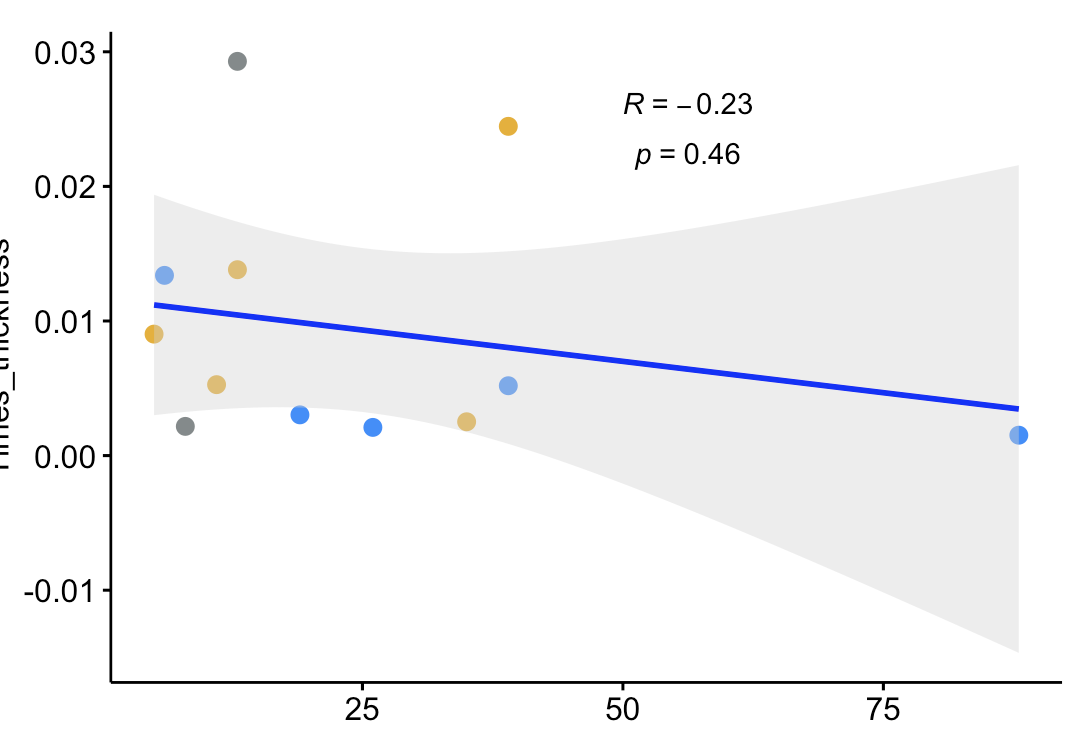

Engagement (total) to stimuli (per 10min)

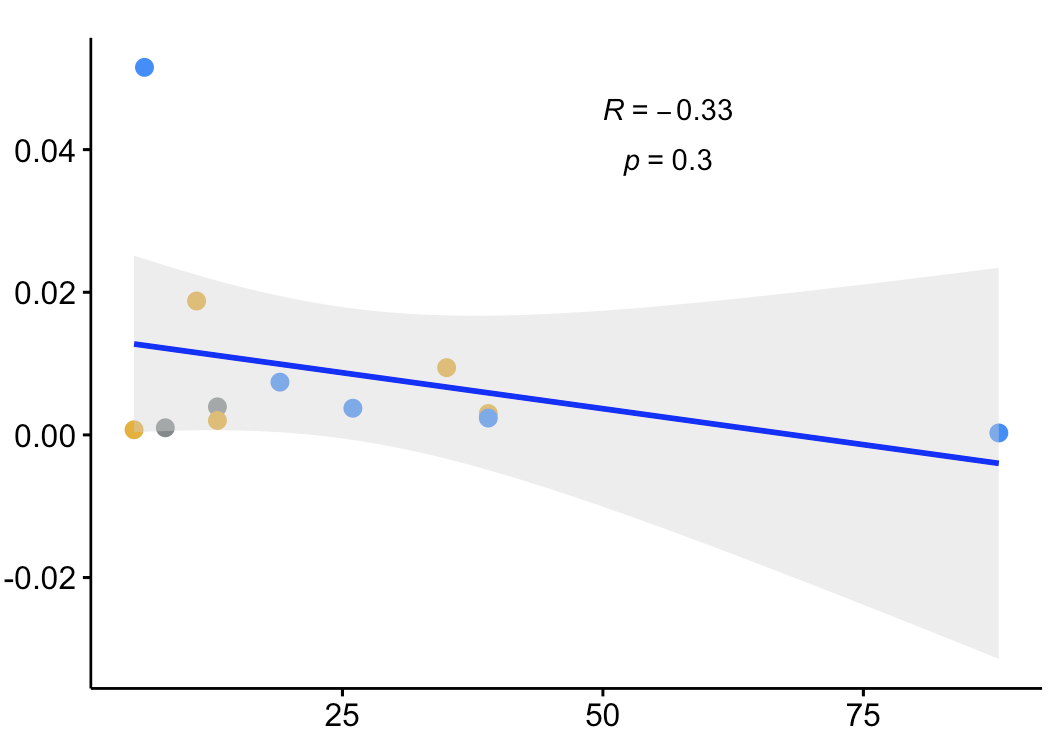

**POA**

**DM**

**Supplemental Figure 2. Relationship between behavioral engagement and neural activity in social decision-making brain regions.** Total engagement with social stimuli during the observational period plotted against pS6/TH colabeled cells across the dorsomedial telencephalon (Dm) and preoptic area (POA). Points represent individual focal males and are colored by social stimulus treatments; lines represent linear regression fits with shaded 95% confidence intervals.

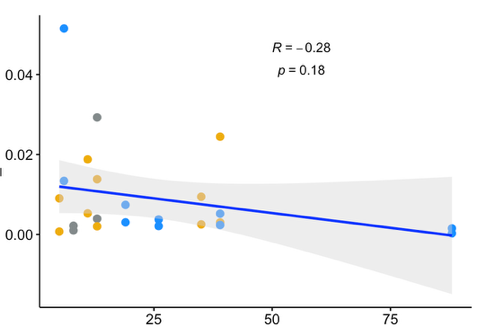

pS6 and TH colabeled cells (averaged by area)

**DM & POA**

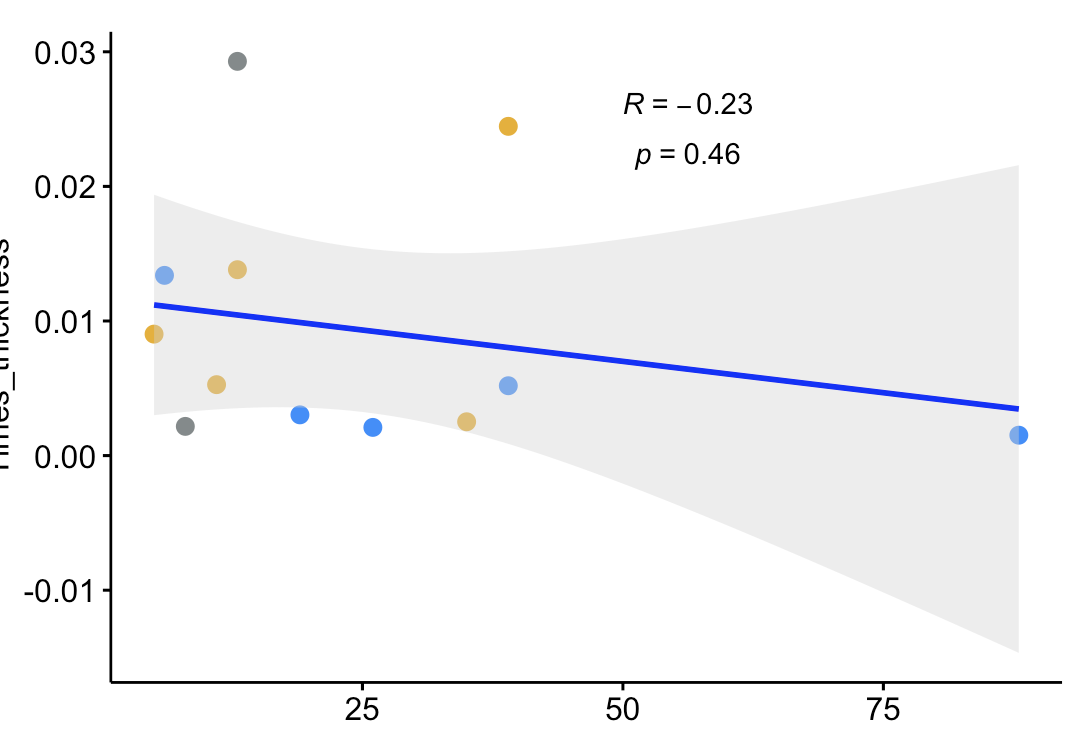

Engagement (total) to stimuli (per 10min)

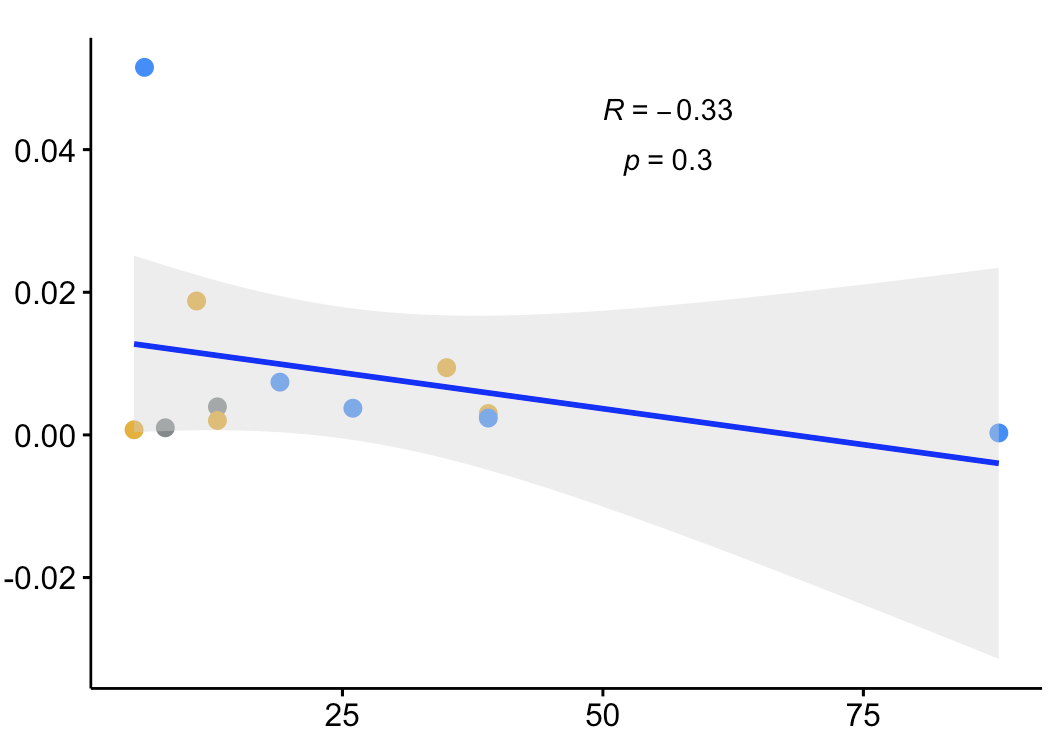

**POA**

**DM**

**Supplemental Figure 2. Examining correlations between total engagement and neural activity across brain regions of interest.**

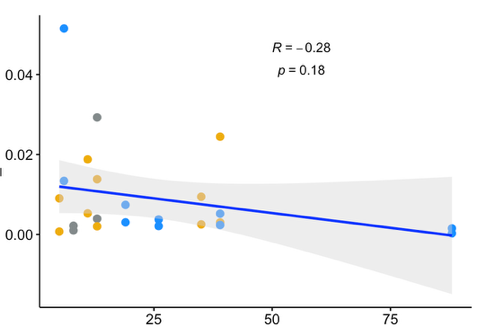

pS6 and TH colabeled cells (averaged by area)

**DM & POA**

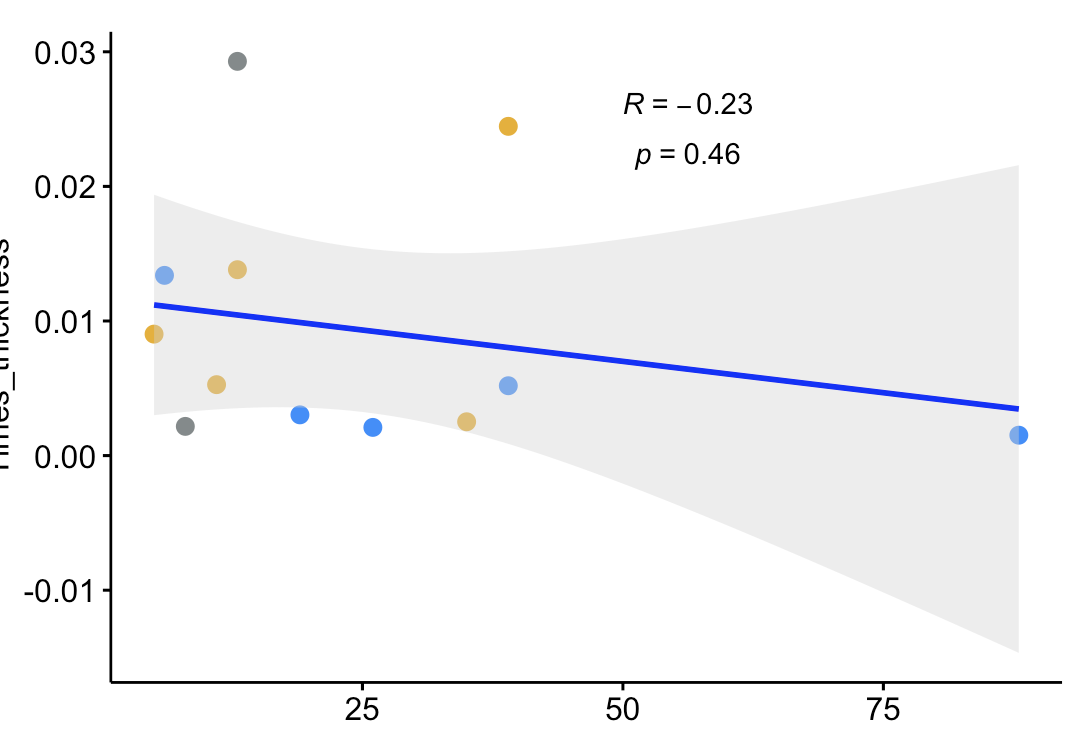

Engagement (total) to stimuli (per 10min)

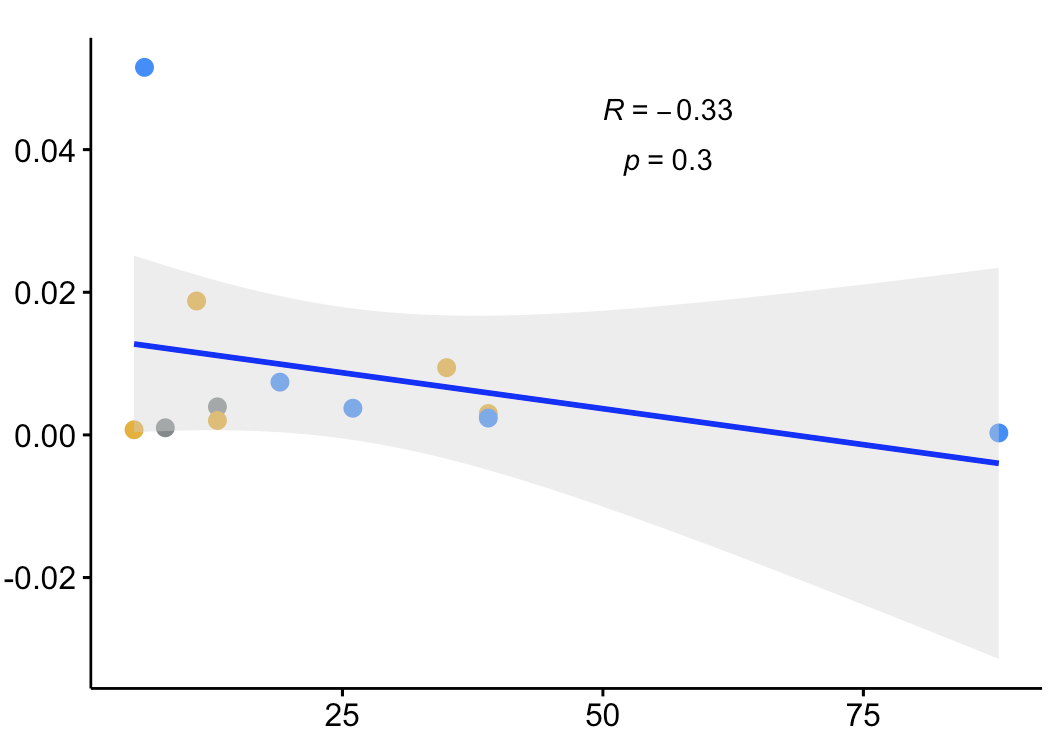

**POA**

**DM**

**Supplemental Figure 2. Examining correlations between total engagement and neural activity across brain regions of interest.**

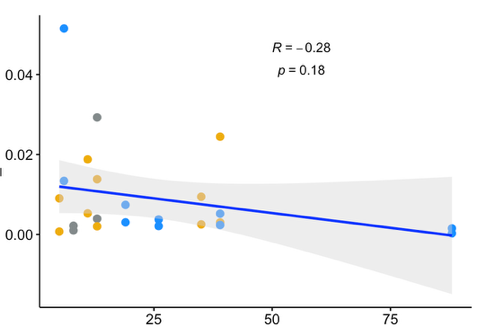

pS6 and TH colabeled cells (averaged by area)

**DM & POA**

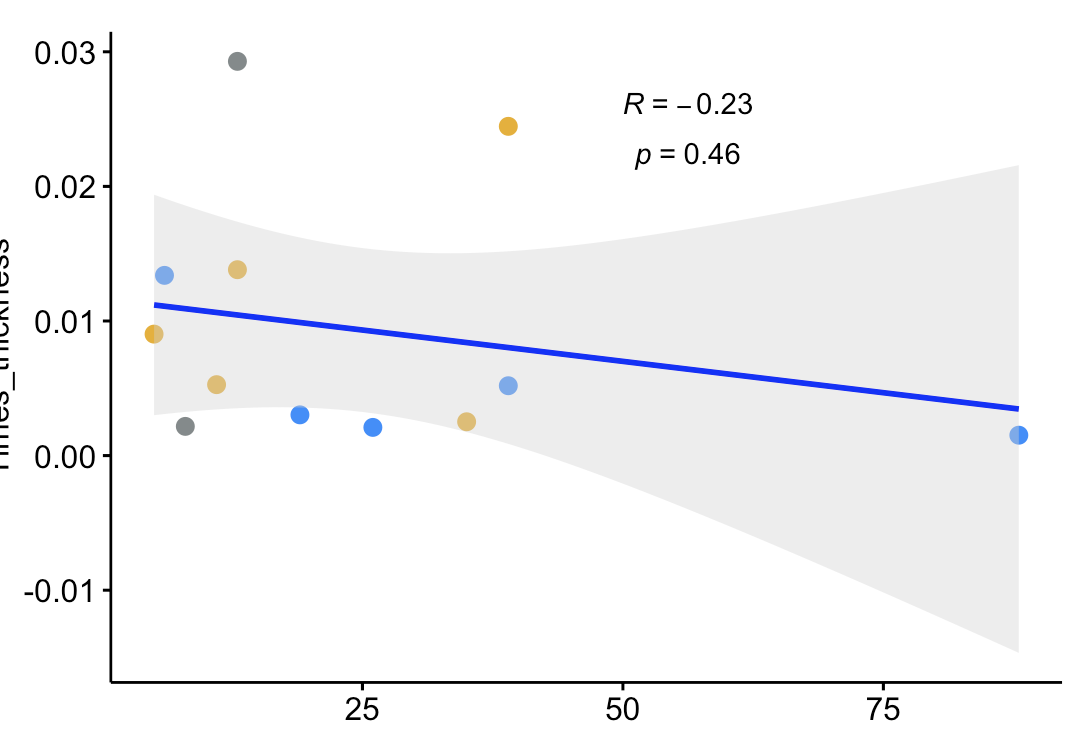

Engagement (total) to stimuli (per 10min)

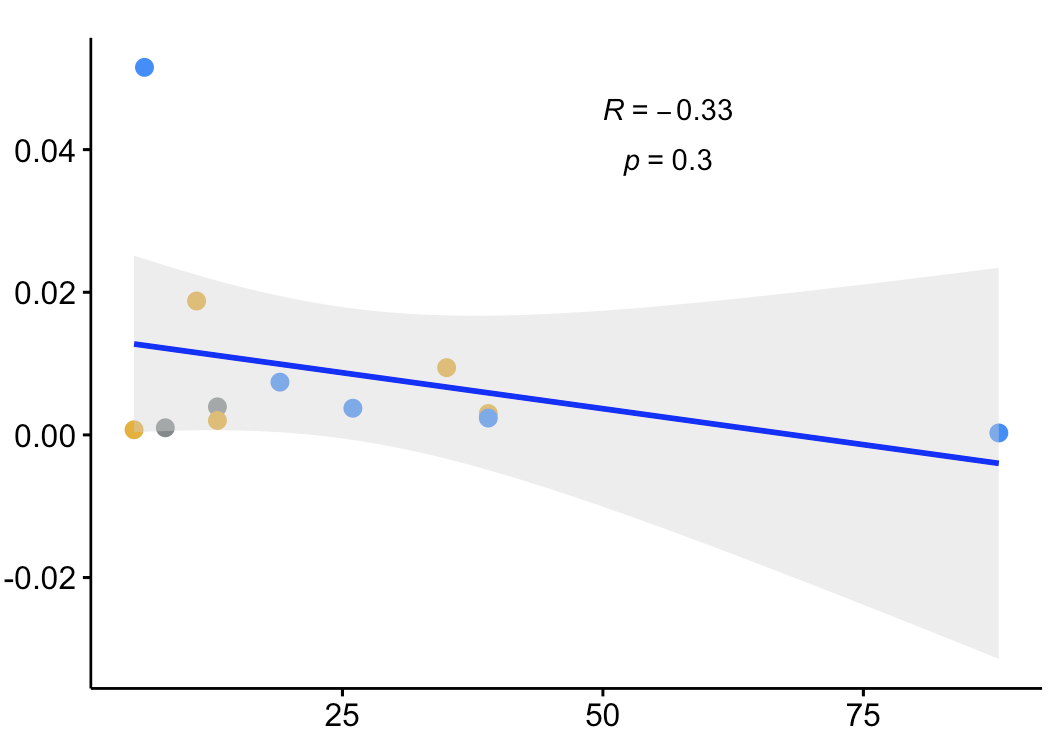

**POA**

**DM**

**Supplemental Figure 2. Examining correlations between total engagement and neural activity across brain regions of interest.**

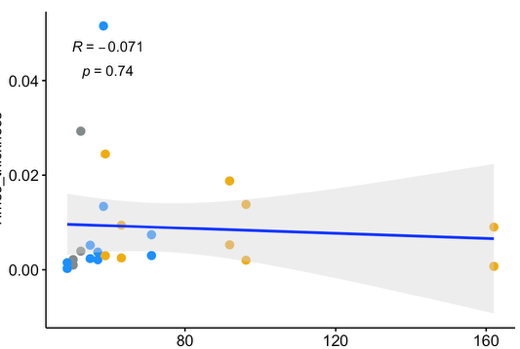

pS6 and TH colabeled cells (averaged by area)

**DM & POA**

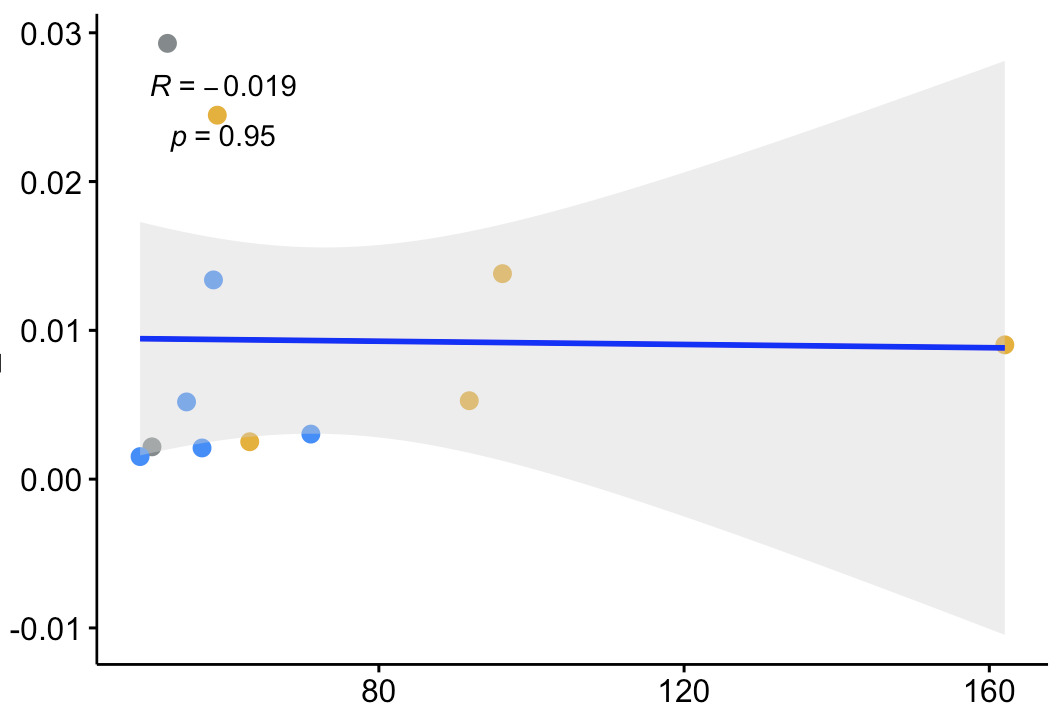

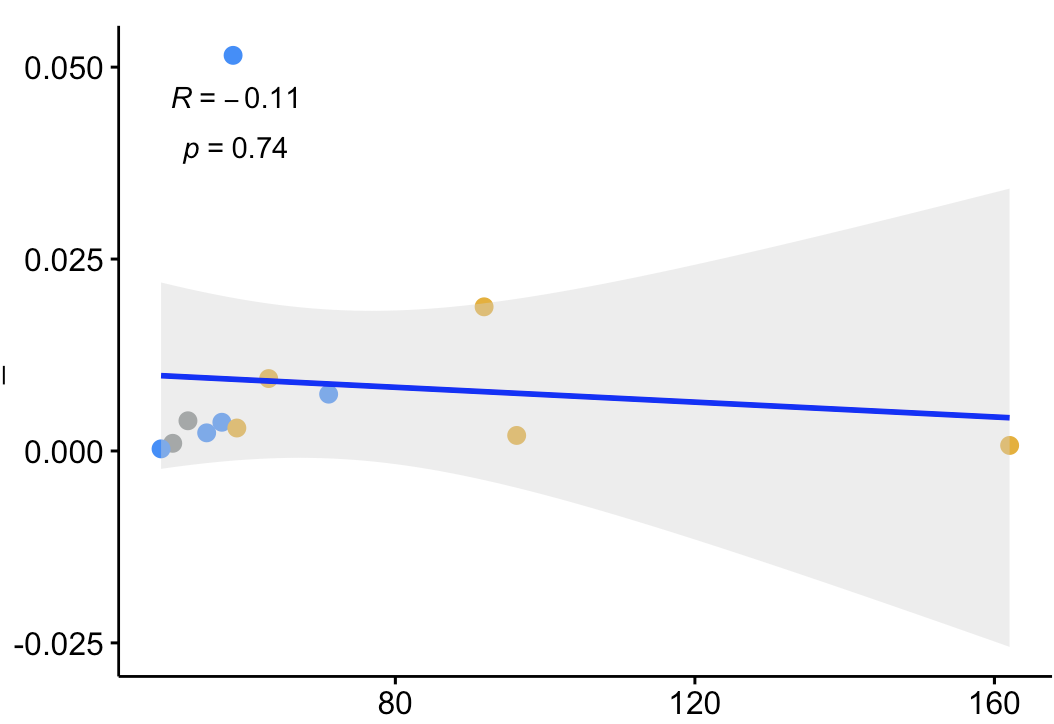

Testosterone (pg/ml)

**POA**

**DM**

**Supplemental Figure 3. Relationship between circulating testosterone and neural activity.** Circulating testosterone levels plotted against pS6/TH colabeled cells across area Dm and POA. Points represent individual focal males and are colored by social stimulus treatment; lines represent linear regression fits with shaded 95% confidence intervals.
